# Hidden heterogeneity and co-occurrence networks of soil prokaryotic communities revealed at the scale of individual soil aggregates

**DOI:** 10.1101/2020.06.24.169037

**Authors:** Márton Szoboszlay, Christoph C. Tebbe

## Abstract

Sequencing PCR-amplified gene fragments from metagenomic DNA is a widely applied method for studying the diversity and dynamics of soil microbial communities. Typically DNA is extracted from 0.25 to 1 g of soil. These amounts, however, neglect the heterogeneity of soil present at the scale of soil aggregates; and thus, ignore a crucial scale for understanding the structure and functionality of soil microbial communities. Here we show with a nitrogen-depleted agricultural soil the impact of reducing the amount of soil used for DNA extraction from 250 mg to approx. 1 mg in order to access spatial information on the prokaryotic community structure as indicated by 16S rRNA-gene amplicon analyses. Furthermore, we demonstrate that individual aggregates from the same soil differ in their prokaryotic communities. The analysis of 16S rRNA gene amplicon sequences from individual soil aggregates allowed us, in contrast to 250 mg soil samples, to construct a co-occurrence network that provides insight into the structure of microbial associations in the studied soil. Two dense clusters were apparent in the network, one dominated by *Thaumarchaeota*, known to be capable of ammonium oxidation at low N concentrations, and the other by *Acidobacteria* subgroup 6 probably representing an oligotrophic lifestyle to obtain energy from SOC. Overall this study demonstrates that DNA obtained from individual soil aggregates provides new insights into how microbial communities are assembled.

## Introduction

One of the major challenges in microbial ecology is to gain a predictive understanding of microbial diversity through elucidating the principles, patterns, and interactions which lead to the assembly of highly diverse microbial communities as found for example in soil (Green, Bohannan, & Whitaker, 2008). To achieve this, it is essential to consider temporal and spatial variation in micro-habitat conditions. In soil microbial ecology, the latter, however, is commonly ignored (Lombard, Prestat, van Elsas, & Simonet, 2011; Vos, Wolf, Jennings, & Kowalchuk, 2013). Instead, large composite samples are favored in order to obtain an overview of microbial diversity at the scale of a plot or a field. This neglects the fine scale heterogeneity of soil structure, and thus loses much information on patterns of community assembly (Thakur et al., 2020).

Soil structure develops as primary particles of different sizes and mineral composition, i.e., clay, silt, and sand, interact with each other, and with organic material to build microaggregates and macroaggregates, that are below or above 250 µm in diameter respectively (Six, Elliott, & Paustian, 2000). Most bacterial cells occur inside aggregates rather than on their surfaces (Ranjard, Poly, et al., 2000) and biogeochemical cycles, which are key ecosystem services driven by an interacting microbial community (Smith et al., 2015), are considered to mainly take place within aggregates (Wilpiszeski et al., 2019). Soil aggregates have been regarded as “massively concurrent evolutionary incubators” (Rillig, Müller, & Lehmann, 2017), or as “microbial villages” (Wilpiszeski et al., 2019), that represent small communities separated by distance and physical barriers and connected only periodically, e.g. during wetting events.

DNA-based methods to assess microbial diversity typically start with extracting 250 mg to 1 g of soil material (Young, Rawlence, Weyrich, & Cooper, 2014). This strategy has been useful to investigate the overall microbial diversity of soils (Fierer & Jackson, 2006; Roesch et al., 2007), its variation across geographical regions (Griffiths et al., 2011; Karimi et al., 2018), or its response to land use change at a continental scale (Szoboszlay, Dohrmann, Poeplau, Don, & Tebbe, 2017). However, in order to understand the processes and interactions taking place within soil microbial communities, it would be rewarding to increase the spatial resolution of the community analysis to individual aggregates and investigate microbial diversity in these spatial entities. Approaching the “calling distance” of microbial interactions (Nunan, 2017), this would increase the likelihood of detecting interacting microbial partners. Correlation networks have increasingly been applied to reveal relationships between microbial community members as detected by PCR amplicon sequence analyses (Banerjee et al., 2016; Barberan, Bates, Casamayor, & Fierer, 2012; Karimi et al., 2020; F. Li, Chen, Zhang, Yin, & Huang, 2017). However, to interpret a positive correlation as a mutualistic and a negative correlation as an antagonistic relationship is only possible for community members sharing the same microhabitat (Weiss et al., 2016). Without distinction of micro-habitats it is hard to assign the presence of taxa to niche exclusion (Faust & Raes, 2012). Analyzing soil DNA extracted from 250 mg to 1 g represents mixed DNA from many microhabitats. In contrast, working with individual soil aggregates, should strongly enhance the ecological significance of soil microbial networks analyses.

The potential impact of the heterogeneous soil constituents on structuring microbial communities at a microscale has already been demonstrated with pooled soil primary particles, where the majority of abundant bacterial and fungal taxa exhibited particular preferences for clay, silt, or sand-sized fractions (Hemkemeyer, Christensen, Tebbe, & Hartmann, 2019; Hemkemeyer, Dohrmann, Christensen, & Tebbe, 2018). Furthermore, comparing pooled samples of micro- and macroaggregates revealed that these two size classes also differ in microbial community composition (Constancias et al., 2014; Davinic et al., 2012; Fox et al., 2018), diversity (Bach, Williams, Hargreaves, Yang, & Hofmockel, 2018; Ivanova et al., 2015) and their response to stress (Ranjard, Nazaret, et al., 2000). However, information on the heterogeneity of microbial communities of individual aggregates within specific aggregate fractions is still lacking. A major limitation of analyzing individual aggregates is the difficulty of obtaining a sufficient quantity of nucleic acids from small amounts of soil for molecular methods. Attempts made so far, therefore, either pooled several aggregates for DNA extraction (Bach et al., 2018; Bailey, Fansler, Stegen, & McCue, 2013; Ivanova et al., 2015), sampled very large aggregates weighing 20 – 70 mg (Kravchenko et al., 2014), or applied whole genome amplification (WGA) (Bailey, McCue, et al., 2013). These solutions, however, do not deliver data on individual aggregates, provide coarse spatial resolution, or generate substantial bias in the results (Direito, Zaura, Little, Ehrenfreund, & Roling, 2014; Wang et al., 2016), respectively. To our knowledge, the only study that reported the bacterial community composition in smaller, i.e. below 3 mm, individual aggregates without applying WGA utilized taxonomic microarrays; a method of relatively low resolution, and focused solely on linking enzyme activity profiles with community structure (Kim, Nunan, Dechesne, Juarez, & Grundmann, 2015). Furthermore, applying molecular methods to small samples that yield very low amounts of nucleic acids require validation to prove the consistent performance of the methods and rule out the possibility of contamination and stochastic effects influencing the results.

The tremendous scientific potential that individual soil aggregate-based microbial community analysis should have for characterizing the heterogeneity of soil microbial communities at a biologically and ecologically more meaningful scale motivated us testing the following hypotheses in this study:

1. Metagenomic DNA of sufficient quantity and quality for PCR-based analyses can be extracted from soil samples in the mg-range, thus representing the scale of macro-aggregates
2. Increasing spatial resolution reveals heterogeneity in soil bacterial and archaeal community structure and abundance
3. A higher heterogeneity seen among small soil samples is not a result of contamination or sub-optimal performance of molecular methods
4. Comparing individual aggregates from the same soil unveils patterns of microbial co-occurrence within the soil microbial community not seen with the commonly used 250 mg soil sample size.

## Materials and methods

### Overview of the experiments

Three experiments were conducted in this study. In the 1^st^ experiment, samples decreasing in size from 250 mg to 1 mg taken from the same soil were subjected to DNA extraction. To address the first two hypotheses, qPCR and high-throughput amplicon sequencing targeting the 16S rRNA gene were conducted to characterize the bacterial, archaeal, and fungal abundance and the prokaryotic diversity in these samples. The 2^nd^ experiment addressed the third hypothesis by comparing 250 mg soil samples and aliquots of a homogenized soil slurry. The volumes of the aliquots were chosen to contain the amount of DNA expected from 1 mg, 5 mg, and 25 mg soil samples. Since all aliquots were taken from the same thoroughly homogenized soil slurry, differences in prokaryotic community structure between these soil homogenate samples must be results of contamination, stochastic effects, or sub-optimal performance of the DNA extraction and PCR. In the 3^rd^ experiment, DNA was extracted from individual aggregates and 250 mg soil samples taken from the same soil to address the fourth hypothesis. All experiments included several control samples to test for the presence of contamination.

### Soil sampling and DNA extraction

The soil used in all experiments was a loam topsoil of a Haplic Chernozem (FAO classification) from the Bad Lauchstädt experimental research station of the Helmholtz Centre for Environmental Research in Germany (51°24’N 11°53’E) (Merbach & Schulz, 2013). It originated form the Static Fertilization Experiment, initiated in the year 1902, and samples were collected in December from a plot without any fertilization since 1903 (treatment NIL)(Ludwig et al., 2011) which was under long term sugar beet – potato – winter wheat – barley rotation. Consequently, the soil was compared to its fertilized variants depleted in nitrogen (Blair, Faulkner, Till, Körschens, & Schulz, 2006). The soil samples had a pH value of 7.1 (in 0.01 M CaCl_2_) and 17.7 mg kg^-1^ organic C. It was sieved (2 mm mesh size) and stored at 4°C until use.

Before sampling, approximately 100 g soil was incubated at room T in the dark for 24 h. The soil was then spread out in a sterile petri dish and samples were taken with sterilized spatulas directly into the bead-beating tubes of the DNA extraction kit. Control samples were included in all experiments. They received no soil but were handled together with and the same way as the soil samples. DNA was extracted with the Quick-DNA Fecal/Soil Microbe Microprep Kit (Zymo Research, Freiburg, Germany) including two 45 s bead beating cycles in an MP FastPrep-24 5G Instrument (MP Biomedicals, Eschwege, Germany) at 6.5 m/s speed with a 300 s break in between. The DNA extracts were eluted in 30 µl elution buffer. All work was done in a biosafety cabinet decontaminated with UV light to minimize the chance of contamination. Measurement of the DNA yield was attempted with Quant-iT PicoGreen dsDNA Assay Kit (Molecular Probes, Life Technologies, Eugene, Or) but accurate quantification was not possible for many of the small samples due to results close to the background fluorescence in the blank controls. In preparation of this study, several DNA extractions methods were tested but were not found suitable. This included the Fast DNA Spin kit for soil (MP Biomedicals, LLC, Illkrich, France) and variations of the phenol-chloroform protocol (Miller, Bryant, Madsen, & Ghiorse, 1999).

In the 1^st^ experiment, five different size-groups of soil samples were collected: 250 mg, 125 mg, 25 mg, 5 mg, and 1 mg, respectively. Eight samples were taken from each size-group along with six control samples. Sample weights from all experiments are listed in Table 1. In the 2^nd^ experiment, ten samples of 250 mg soil and six control samples were taken. Eight of the soil samples were processed normally in the DNA extraction, while for the other two, the DNA extraction was interrupted after the centrifugation following the bead beating. By this point, the samples had been turned into homogenized slurry by the bead beating, and the soil debris had been separated from the supernatant that contained the metagenomic DNA. The supernatant from the two samples were pooled and homogenized by vortexing. The mass of the resulting suspension was 1 112 mg and it originated from 497 mg soil in total. Accordingly, a 55.9 mg aliquot of this suspension contained the amount of DNA extractable from 25 mg soil, an 11.2 mg aliquot the amount from 5 mg soil, and a 2.2 mg aliquot the amount from 1 mg soil. Eight aliquots from each of these sizes, hereafter referred to as 25 mg, 5 mg, and 1 mg soil homogenate samples, were taken and each mixed with 350 µl bashing bead buffer from the DNA extraction kit to continue the DNA extraction. In the 3^rd^ experiment, 37 individual soil aggregates, weighing 5.3 mg on average and similar in size (ca. 2 mm), were taken for DNA extraction along with 35 samples of 250 mg soil and nine control samples.

**Table 1.** Number of samples and soil weights within each sample category.

| Soil weight class or sample type | Sample weight [mg] ± SD | Number of soil samples |
| --- | --- | --- |
| *1<sup>st</sup> Experiment* |  |  |
| 250 mg | 251 ± 1 | 8 |
| 125 mg | 125 ± 1 | 8 |
| 25 mg | 25.1 ± 0.4 | 8 |
| 5 mg | 4.9 ± 0.2 | 8 |
| 1 mg | 1.1 ± 0.2 | 8 |
| Control, no soil |  | 6 |
| *2<sup>nd</sup> Experiment* |  |  |
| 250 mg | 252 ± 3 | 8 |
| 25 mg soil homogenate |  | 8 |
| 5 mg soil homogenate |  | 8 |
| 1 mg soil homogenate |  | 8 |
| Control, no soil |  | 6 |
| *3<sup>rd</sup> Experiment* |  |  |
| 250 mg | 251 ± 0 | 35 |
| Soil aggregate | 5.3 ± 1.6 | 37 |
| Control |  | 9 |

### Abundances of microbial groups assessed by qPCR

The abundance of *Bacteria, Archaea*, and *Fungi* were assessed by qPCR targeting the 16S rRNA gene and the ITS region as previously described (Hemkemeyer, Christensen, Martens, & Tebbe, 2015). Archaeal and fungal abundance was only investigated in the 1^st^ experiment. Reactions were run in a Bio-Rad CFX96 real-time PCR system in duplicates from different dilutions of the DNA extracts. In case of the 250 mg and 125 mg soil samples, 50- and 100-fold dilutions were taken; from the 25 mg samples 10- and 20-fold dilutions; and from the 5 mg, 1 mg, individual aggregate, and control samples undiluted DNA extracts and two-fold dilutions were used. PCR efficiencies were 95.9 – 104.6% for the bacterial 16S rRNA gene, 94.2 – 96.2% for the archaeal 16S rRNA gene, and 83.3 – 84.3% for the fungal ITS with R^2^ ≥ 0.995 in all cases. Results were compared with Tukey’s HSD tests, or Welch’s t-test in case of the data from the 3^rd^ experiment. The analysis was carried out in R 3.4.4 (www.r-project.org). One of the 250 mg samples from the 1^st^ experiment yielded a magnitude higher copy number in the fungal ITS qPCR assay than the others. It was deemed to be an outlier and excluded from the analysis.

### High-throughput sequencing of 16S rRNA gene amplicons and data processing

To characterize the bacterial and archaeal communities in the samples, DNA extracts were subjected to high-throughput amplicon sequencing of the V4 region of the 16S rRNA gene following the protocol of Kozich *et al*. (Kozich, Westcott, Baxter, Highlander, & Schloss, 2013) with primers updated to match the modified 515f and 806r sequences according to Walters *et al*. (Walters et al., 2016). In case of the small soil samples and control samples, due to the low DNA yield, 10 µl DNA extract was used as template in the PCRs. Paired-end sequencing was done on Illumina MiSeq instruments at StarSEQ (Mainz, Germany). Samples from the same experiment were sequenced in the same run. All data is available at the European Nucleotide Archive under the accession numbers PRJEB36881 (1^st^ experiment), PRJEB36883 (2^nd^ experiment), and PRJEB36887 (3^rd^ experiment).

The sequencing data from the three experiments were analyzed separately. Raw reads were processed with the dada2 (version 1.6.0) pipeline (Callahan et al., 2016) in R 3.4.4. Forward and reverse reads were truncated at positions 240 and 90, respectively. Reads with any ambiguous bases were discarded as well as forward reads with over two and reverse reads with over one expected error. The data from the 2^nd^ experiment had higher quality allowing the reverse reads to be truncated at position 130 and keeping those with two or less expected errors. Error models were constructed from 10^6^ randomly selected reads. Sequence variants (SVs) were inferred using the pool option. Forward and reverse SVs were merged trimming overhangs, and the removeBimeraDenovo function was employed to detect chimeras. The SVs were classified according to the SILVA reference version 132 (Pruesse et al., 2007) accepting only results with ≥70% bootstrap support. SVs shorter than 220 nt or longer than 275 nt, or identified as chimeric, mitochondrial, or chloroplast sequences, or not classified into *Bacteria* or *Archaea* were deleted. Good’s index was calculated to estimate the coverage of the SVs. SVs with ≥0.1% relative abundance in any of the control samples of an experiment were regarded as potentially contaminant and removed from the dataset of that experiment.

### Analysis of the sequencing results

Principal component analysis (PCA) plots were created in R using the rda function of the vegan package version 2.5-2 (Oksanen et al., 2018). To decrease the sparsity of the data, SVs not reaching 0.1% relative abundance in any of the samples included in the PCA were removed. Zeroes were replaced with the count zero multiplicative (CZM) method using the zCompositions package version 1.1.1 (Palarea-Albaladejo & Martin-Fernandez, 2015) and centered log-ratio (CLR) transformation was applied to the data to correct for compositional effects and differences in sequencing depth (Gloor, Wu, Pawlowsky-Glahn, & Egozcue, 2016).

Aitchison distances between the samples were calculated as Euclidean distances in the CLR transformed dataset (Gloor, Macklaim, Pawlowsky-Glahn, & Egozcue, 2017). SVs that didn’t have at least 0.1% relative abundance in any of the compared samples were removed and zeroes were replaced with the CZM method to allow CLR transformation before calculating Euclidean distances with the ‘vegdist’ function of the vegan package. Results were compared with Tukey’s HSD tests.

Plots illustrating the prevalence of abundant SVs among the soil samples were prepared in Cytoscape 3.7.1 (www.cytoscape.org). CoNet 1.1.1 beta (K Faust & Raes, 2016) in Cytoscape was used to construct co-occurrence networks from the data from the 3^rd^ experiment. To limit the number of parallel significance tests and the sparsity of the data, only SVs with ≥0.2% relative abundance in at least one of the samples were included. Separate networks were constructed for the 250 mg soil samples and the soil aggregates. However, the selection of SVs was done on the joint data matrix to ensure that both networks include the same SVs. The data was relativized to the total sequence count of each sample. Pearson and Spearman correlations, mutual information (jsl setting), and Bray-Curtis and Kullback-Leibler (with a pseudo count of 10^−8^) dissimilarities were calculated and the 1 000 highest and 1 000 lowest scoring edges from each of the five metrics were kept. The ReBoot method (Faust et al., 2012), which mitigates compositional effects, was used to assess the significance of the edges based on 1 000 permutations with renormalization and 1 000 bootstrap iterations. In the network of the soil aggregates, edges with scores below the 2.5^th^ and over the 97.5^th^ percentile of the bootstrap distribution or not supported by at least three of the five metrics were considered unstable and removed. Brown’s method of p-value merging was applied followed by Benjamini-Hochberg correction. Only edges with q ≤ 0.05 were included in the final network. In the network of the 250 mg samples, unstable edges were not removed and the Benjamini-Hochberg correction was not applied as otherwise no edges were retained. The networks were visualized in Cytoscape using the compound spring embedder layout. Topological parameters were calculated with NetworkAnalyzer version 2.7 (Assenov, Ramirez, Schelhorn, Lengauer, & Albrecht, 2008).

## Results

### Microbial DNA can be extracted from soil samples in the mg-range

DNA could be extracted from soil samples as little as 0.87 mg as well as from intact soil aggregates. In all cases, the extracted DNA was sufficient for 16S rRNA gene amplicon sequencing and the qPCR assays. Accurate quantification of the DNA yield with PicoGreen assays was not possible because the fluorescence readings from many of the small samples were close to the background fluorescence in the blank controls.

To assess whether the DNA extraction could recover microbial DNA with similar efficiency from small quantities of soil as from 250 mg samples, estimates of the abundances of *Bacteria, Archaea*, and *Fungi* in one gram of soil were calculated from the qPCR results (Figure 1; Supplementary data file #1). Similar estimates were obtained from the 250 mg and 5 mg soil samples from the 1^st^ experiment. Estimates from the 1 mg samples tended to be lower but were not significantly different. In contrast, the estimates of fungal abundance in a gram of soil were significantly lower from the 1 and 5 mg than from the 250 mg samples. Estimates of bacterial abundance obtained from the 25 mg and 125 mg samples and archaeal abundance from the 25 mg samples were significantly higher than from the 250 mg samples. Bacterial abundance estimates from the single aggregates and 250 mg soil samples of the 3^rd^ experiment covered the same range (Figure S1), but the mean of the estimates from the single aggregates (2.27 × 10^9^ copies/g soil) was lower (p < 0.001) than from the 250 mg samples (4.69 × 10^9^ copies/g soil).

**Figure 1.**
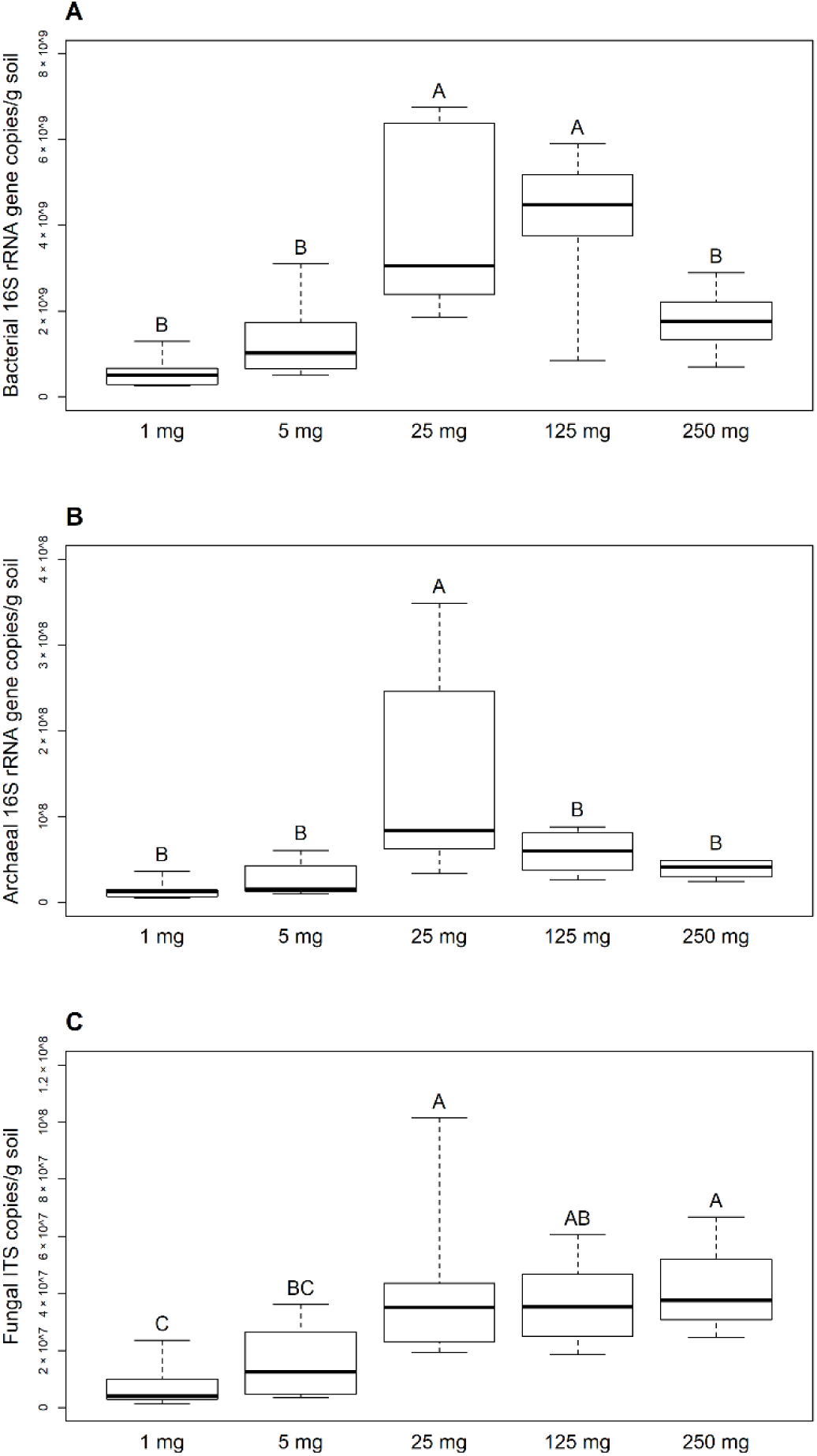
Estimates of (A) bacterial, (B) archaeal, and (C) fungal abundance in a gram of soil based on qPCR from the samples from the 1^st^ experiment (gene copy numbers per g of soil wet weight). One of the 250 mg soil samples was an outlier in the fungal ITS qPCR results and is not included in the plot. Sample groups not labelled with the same letter were significantly different in Tukey’s HSD tests. Thick lines indicate the median values, the upper and lower hinges the 75^th^ and 25^th^ percentile, whiskers extend to the data extremes.

The control samples amplified in the qPCR assay targeting the bacterial 16S rRNA gene but yielded only 3 to 356 copies per µl DNA extract. In comparison, the 1 mg soil samples had 8 777 to 45 534 copies per µl DNA extract (Supplementary file 1). The control samples from the 1^st^ experiment had 0 to 13 fungal ITS copies per µl DNA extract and none of them showed amplification in the archaeal 16S rRNA gene qPCR assays.

### Removing potentially contaminant SVs from the sequencing results

It was possible to generate sequencing results from all control samples (the complete dataset with the taxonomic classification of the SVs is in Supplementary file 2). Good’s coverage index of the SVs was over 0.993 in all of them indicating that almost their complete prokaryotic community was captured by the sequencing (Table S1). They were very similar to each other in their prokaryotic community structures but very different from the soil samples (Figure S2). Every SV that reached 0.1% relative abundance in any of the control samples of an experiment was considered as a potential contaminant. There were 450, 77, and 591 such SVs in the datasets of the 1^st^, 2^nd^, and 3^rd^ experiments respectively. In the data from the 1^st^ experiment, these SVs together covered 98.9 – 99.8% of the sequences obtained from the control samples and 2.5 – 6.6% of the sequences from the soil samples. In the 2^nd^ and 3^rd^ experiments, 92.5 – 99.5% and 98.6 – 99.6% of the sequences from the control samples, and 2.9 – 17.3% and 5.9 – 9.4% of the sequences from the soil samples, respectively, were covered by the potentially contaminant SVs. To mitigate the effect of contamination on the results, the potentially contaminant SVs were deleted from the data matrices before further analysis.

### Increasing spatial resolution reveals heterogeneity in soil bacterial and archaeal community structure but not in their abundance

The yield of high-quality 16S rRNA gene amplicon sequences was somewhat lower from the 1 mg samples than from the 5 mg and 25 mg samples in the 1^st^ experiment (Figure 2A). Apart from this, however, the sequencing yield did not differ between the sample groups within any of the experiments. Thus, it is possible to compare the number of SVs detected in the samples without rarefying the data. The number of SVs in the 1^st^ experiment was not significantly different between the 250 mg, 125 mg, and 25 mg samples, but decreased significantly in the 5 mg and even further among the 1 mg samples (Figure 2B). An opposite trend was apparent in the Good’s coverage index (Table S1). Similarly, significantly lower numbers of SVs were detected in the single aggregates than in the 250 mg soil samples of the 3^rd^ experiment.

**Figure 2.**
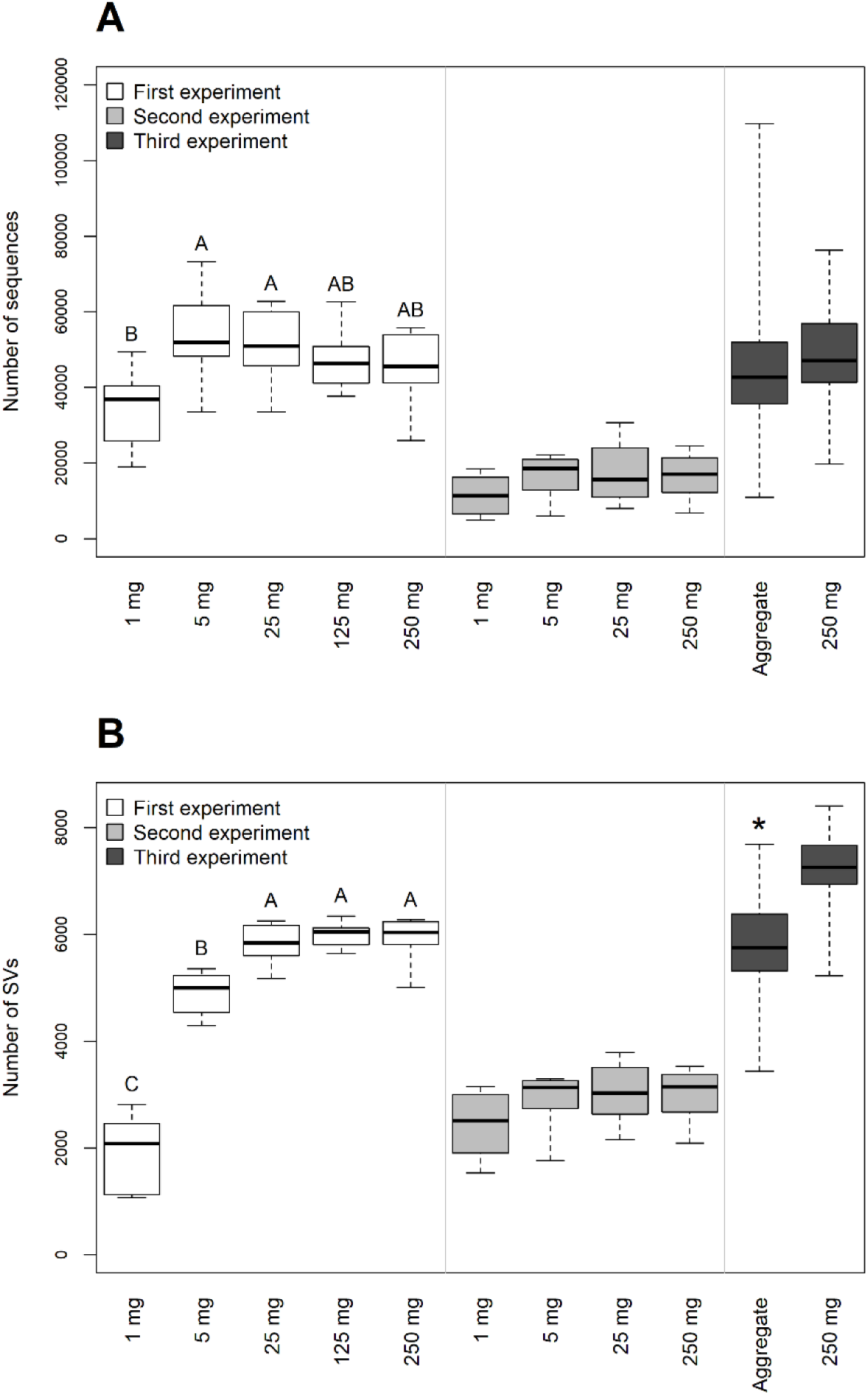
Number of (A) sequences and (B) sequence variants (SVs) in the samples after the removal of potentially contaminant SVs. Thick lines indicate the median values, the upper and lower hinges the 75^th^ and 25^th^ percentile, whiskers extend to the data extremes. Letters indicate significant differences between sample groups of the 1^st^ experiment according to Tukey’s HSD tests. * indicates significant difference based on Welch’s t-test between the aggregate and the 250 mg soil samples of the 3^rd^ experiment.

Principal component analysis (PCA) from the sequencing results from the 1^st^ experiment arranged all 250 mg and 125 mg samples, and most 25 mg samples into a single, tight group, indicating high similarity in their prokaryotic community structures (Figure 3A). In contrast, samples from the 5 mg and more so from the 1 mg categories, showed higher heterogeneity in community structure. Similarly, PCA indicated heterogeneity between individual aggregates that was not seen among the 250 mg samples in the 3^rd^ experiment (Figure 3B).

**Figure 3.**
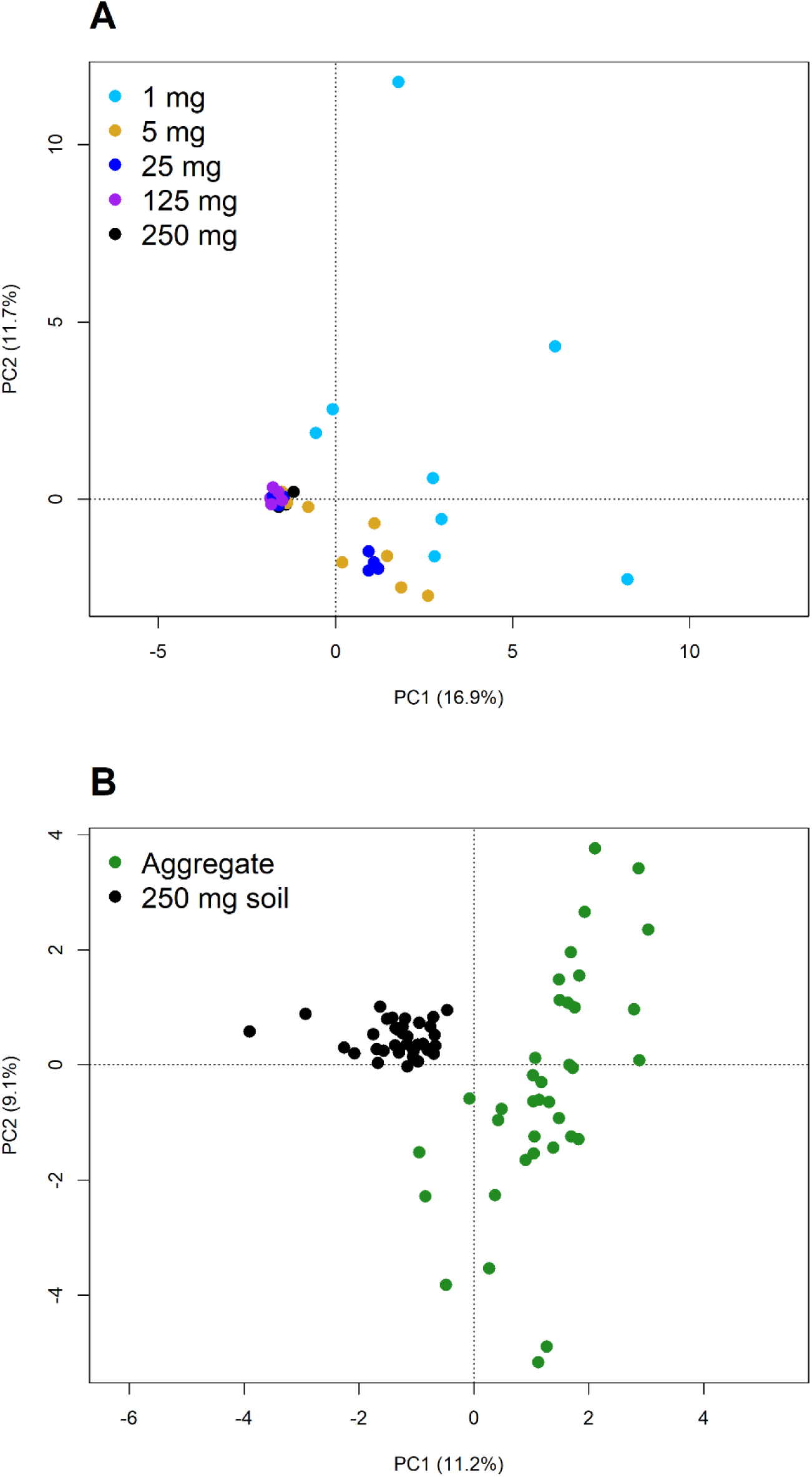
Principle component analyses (PCA) plots from the 16S rRNA gene sequencing data from the 1^st^ (A) and 3^rd^ (B) experiments.

The abundant SVs (≥ 0.1% relative abundance in at least one of the samples) in the 250 mg, 125 mg, and 25 mg samples from the 1^st^ experiment were almost all detectable in each individual sample, showing that the composition of the soil prokaryotic community appears very uniform when investigated at such a coarse spatial resolution (Figure 4). In contrast, 172 of the abundant SVs detected in the 1 mg samples were unique to just one or two of these samples. Among these SVs, representatives of *Planctomycetes, Proteobacteria*, and *Acidobacteria* were especially numerous, while *Thaumarchaeota* and *Actinobacteria* were dominant among the SVs present in all samples. The 5 mg samples represented a level of spatial resolution at which some heterogeneity in the prevalence of the abundant SVs was already apparent with 22 of them detectable in two or only in a single sample.

**Figure 4.**
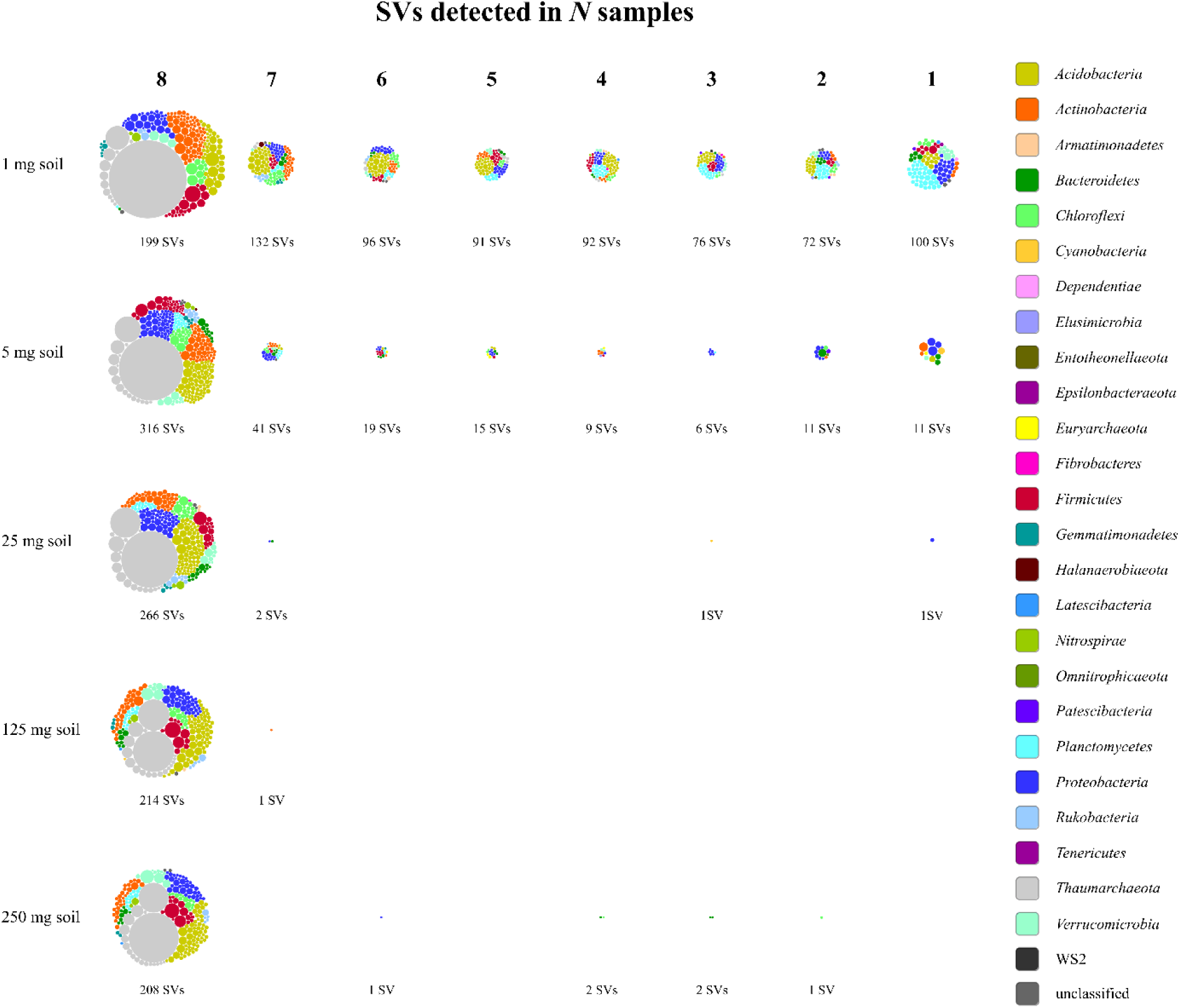
SVs arranged according to in how many of the 1 mg, 5 mg, 25 mg, 125 mg, or 250 mg samples from the 1^st^ experiment they were detected. Only SVs that reached at least 0.1% relative abundance in any of the samples are included. Each node represents one SV colored based on its phylum-level classification and sized according to its average relative abundance across all samples excluding those in which it was not detected.

In total, 5 620 SVs were detected in the eight 1 mg soil samples of the 1^st^ experiment (Supplementary file 2). Of these, 4 764 (85%) were also present in at least half of the 250 mg samples. The remaining 856 SVs had low relative abundance in the 1 mg samples with only 59 reaching more than 0.1% relative abundance in any of them. The 5 mg samples together contained 8 010 SVs, of which 6 443 (80%) were also detectable in at least half of the 250 mg samples. Of the remaining 1 567 SVs, only 26 reached more than 0.1% relative abundance in any of the 5 mg samples.

The qPCR results did not confirm our hypothesis that increasing spatial resolution would reveal heterogeneity in microbial abundance. Bacterial and archaeal 16S rRNA gene, and fungal ITS copy numbers did not show a larger variation among the 1 and 5 mg samples than between the 250 mg samples from the 1^st^ experiment (Figure 1). Similarly, in the 3^rd^ experiment, bacterial abundance did not vary more in the single aggregates than in the 250 mg samples (Figure S1).

### Impact of stochastic effects and inconsistent performance of the methods

The soil homogenate samples from the 2^nd^ experiment served to test the influence of stochastic effects and sub-optimal performance of the DNA extraction and PCR when extracting small amounts of soil. The estimated bacterial abundance in a gram of soil based on the qPCR results was in general higher in the samples from the 2^nd^ experiment but followed the same pattern as in the samples from the 1^st^ experiment with no differences between the 1 mg, 5 mg, and 250 mg samples but significantly higher values in the 25 mg samples (Figure S3). The small soil homogenate samples did not show the degree of heterogeneity in prokaryotic community structure we observed among the small soil samples of the 1^st^ experiment. The abundant SVs (≥ 0.1% relative abundance in at least one sample) in the 25 mg soil homogenate samples were all detectable in at least five of the eight replicates. Out of the 354 abundant SVs in the 5 mg soil homogenate samples, one was present in only three of the samples but the others were detectable in at least six. The 1 mg soil homogenate samples harbored 446 abundant SVs. None of them was unique to a single sample and 442 were present in five or more of the eight samples. The Aitchison distances of community structure were much higher among the 5 mg, and especially among the 1 mg soil samples of the 1^st^ experiment compared to the distances between the 250 mg soil samples (Figure 5). In contrast, the distances between 5 mg or 25 mg soil homogenate samples of the 2^nd^ experiment were very similar to the distances among the 250 mg soil samples indicating no difference in the heterogeneity of prokaryotic community structure between these sample groups. The distances between the 1 mg soil homogenate samples were only slightly increased.

**Figure 5.**
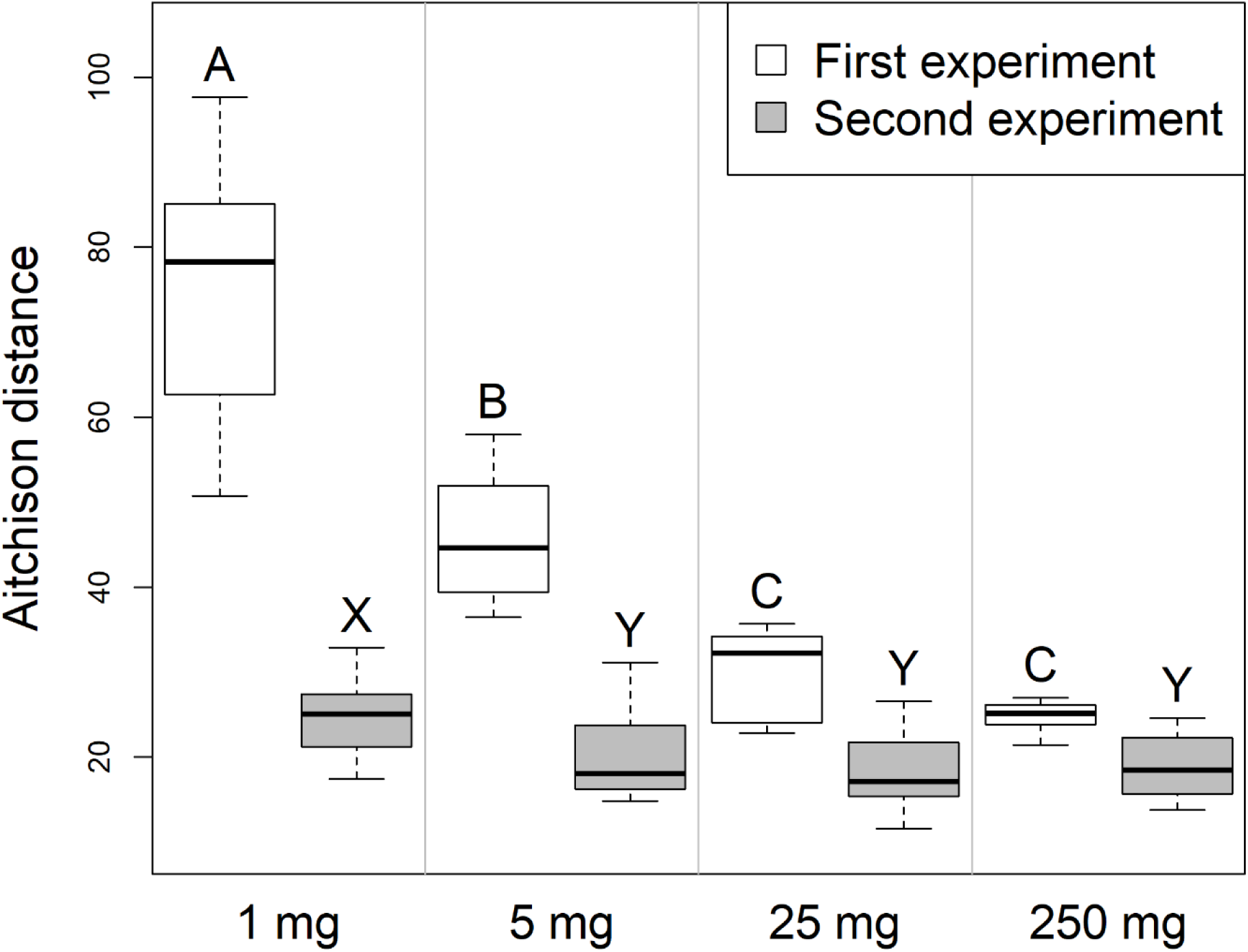
Aitchison distances in the bacterial and archaeal community structure (16SrRNA gene amplicons) within sample groups from the 1^st^ and 2^nd^ experiments. Thick lines indicate the median values, the upper and lower hinges the 75^th^ and 25^th^ percentile, whiskers extend to the data extremes. Sample groups from the same experiment not labelled with the same letter were significantly different in Tukey’s HSD tests.

### Bacterial and archaeal co-occurrence patterns in 250 mg soil samples and aggregates

Networks of prokaryotic co-occurrence were constructed using the 272 SVs that reached ≥0.2% relative abundance in at least one of the samples of the 3^rd^ experiment. No network was obtained from the 250 mg samples unless the removal of unstable edges and the correction of the p-values for multiple testing were skipped. The resulting network has 78 edges between 35 nodes (Figure 6A). Thus, this spatial resolution revealed only a small number of putative associations many of which are likely false discoveries. In contrast, a network of 137 edges and 67 nodes (with the removal of unstable edges and control of the false discovery rate) was obtained from the individual soil aggregates (Figure 6B). A total of 54 of the nodes are part of a connected component in which there are three nodes with high betweenness centrality: SV15 (*Verrucomicrobia*, candidatus *Udaeobacter*), SV31 (*Actinobacteria*), and SV36 (*Acidobacteria* subgroup 6). Their relative abundance in the aggregates were 0.46 ± 0.16%, 0.33 ± 0.13%, and 0.30 ± 0.11%, respectively. These SVs potentially serve a keystone function by connecting two clusters in the network. One of the clusters contains several SVs of *Thaumarchaeota, Verrucomicrobia*, and *Actinobacteria*. The other is dominated by *Acidobacteria* subgroup 6. The hub of the latter cluster is SV58 (*Acidobacteria* subgroup 6) with a relative abundance of 0.23 ± 0.16% that has the highest degree in the network being connected to 19 nodes. SV399 (*Chloroflexi*) (0.06 ± 0.05%) and SV123 (*Acidobacteria* subgroup 6) (0.13 ± 0.06%) are linked with negative associations to several members of this cluster.

**Figure 6.**
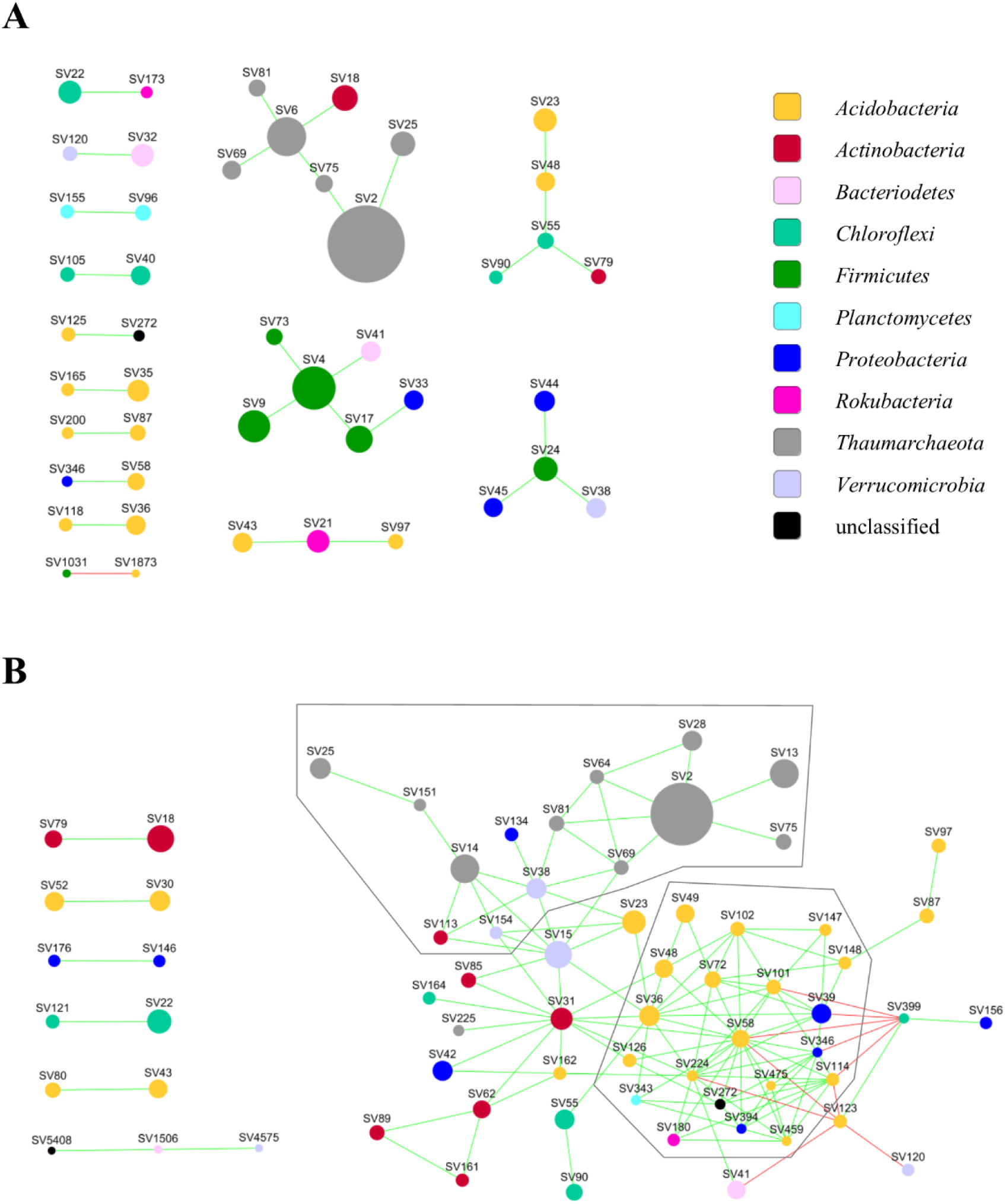
Co-occurrence networks from the (A) 250 mg samples and (B) the single aggregates from the 3^rd^ experiment. The frames mark the two clusters discussed in the text. It should be noted that in (A) unstable edges were *not* removed and the Benjamini-Hochberg correction for multiple comparisons was not applied.

## Discussion

The large disparity between the scale in which the soil microbiota is usually studied with molecular methods (0.25 to 1 g of soil) and the distance over which microbial interactions occur, impede the detection of interacting partners (Nunan, 2017). In order to gain information on the soil microbial diversity at an increased spatial resolution that considers soil structure, in this study we reduced the amount of soil used for DNA extraction from 250 to 1 mg and also extracted individual soil aggregates. Bacterial and archaeal DNA was recovered with not a significantly different efficiency from the 250 mg and the 5 mg and 1 mg samples as shown by the qPCR results. This was not true for fungal DNA. Either the DNA-extraction kit was not efficient in isolating fungal DNA from samples below 25 mg, or fungi may preferentially colonize larger soil aggregates. Our results show that the DNA extraction kit was most efficient in recovering bacterial and archaeal DNA from 25 to 125 mg soil, although the variation in the yield of 16S rRNA gene copies was large among these samples. Since this increased variation was apparent among the 25 mg soil homogenate samples of the 2^nd^ experiment as well, it is not an indication of an uneven distribution of bacterial cells at the scale of 25 mg samples but must be due to this particular DNA extraction method not working with consistent efficiency with this amount of soil.

As a consequence of sampling small amounts of soil, the DNA extracts had low template concentrations for the subsequent PCR analyses. Thereby we had to anticipate a high risk of contamination affecting the results (Weiss et al., 2014). In fact, quantifiable amounts of *Bacteria* and *Fungi*, but not *Archaea*, were detected in the control samples without soil. However, they reached no more than 4% of the number of bacterial rRNA genes copies in the smallest soil samples, and thus, the influence of contamination on our results is negligible. The bacterial community found in the control samples was clearly distinct from the soil communities suggesting that the contamination originated from the reagents of the DNA extraction and sequencing library preparation rather than cross-contamination between samples (Glassing, Dowd, Galandiuk, Davis, & Chiodini, 2016; Salter et al., 2014). Another concern of working with small samples is that molecular methods applied to such low amounts of template may perform inconsistently leading to artificial variation in the results. The 2^nd^ experiment showed that the DNA extraction, PCR, and sequencing did not artificially generate more variation in the results from 5 mg samples than the variation present among the 250 mg samples and the 1 mg samples showed only a slightly higher variation. Therefore, the large heterogeneity in prokaryotic community composition and structure among the 1 mg and 5 mg soil samples from the 1^st^ experiment was not caused by stochastic effects or PCR bias.

The samples of 25 mg up to 250 mg of soil were close to identical in prokaryotic community composition, thus they provide a good representation of the overall prokaryotic diversity of our soil. This is also indicated by the fact that increasing the amount of soil extracted up to 25 mg increased the number of SVs detected in the samples, but larger soil samples did not yield more SVs. Thus, the 25 mg samples already had a good coverage of the total prokaryotic community. In contrast, the 1 mg and 5 mg samples and single aggregates were heterogeneous in community structure. We found that while small soil samples could recover some SVs not necessarily detected with the conventionally used 250 mg samples; these SVs were typically low in abundance. Very few exceeded the relative abundance threshold we applied to control the sparsity of the data in our analysis of community structure. Therefore, the large heterogeneity of prokaryotic community structure we observed among the small samples was not because they would have enabled the detection of more SVs. Instead, it appears that they contained different subsets of the total community present in the 25 to 250 mg samples. This could be explained by the fact that the smaller samples contain fewer micro-habitats, each of which harbors a local community of fewer species (Leibold et al., 2004). Interestingly, the 1 mg and 5 mg samples did not significantly differ in the abundance of *Bacteria, Archaea*, and *Fungi* compared to the 250 mg samples. Similarly, the variation in bacterial abundance found with the individual aggregates was not different from the 250 mg samples. Microbial abundance in soil has a patchy distribution at the scale of a few micrometers (Nunan, Wu, Young, Crawford, & Ritz, 2003) but, for the soil of this study, apparently not at the scale of macroaggregates or 1 to 5 mg samples.

Network analyses based on microbial co-occurrence have been applied to soil samples as large as 10 grams (Khan et al., 2019) and is typically used with 250 mg – 1 g samples (Barberan et al., 2012; Karimi et al., 2020). In this study, however, we could not detect stable and significant associations between SVs from 35 samples of 250 mg soil. These samples were taken from the same well-mixed batch of soil and were very similar in their prokaryotic community composition. It is likely, that each 250 mg sample gave a good representation of the overall prokaryotic diversity in our soil, in which case, the variation of the relative abundance of SVs in these samples were mostly random. Random patterns should not yield stable and significant associations in network analysis. The value of utilizing much smaller samples is shown by our result that the 1 and 5 mg samples contained subsets of the total soil microbial diversity captured by the 250 mg samples, and with increasing spatial resolution the heterogeneity in the bacterial and archaeal community structure increased among the samples. This results in detectable co-occurrence patterns. Furthermore, the smaller spatial scale increases the likelihood that the observed co-occurrences actually indicate interactions (Cordero & Datta, 2016).

From 37 soil aggregates, we obtained a complex network of bacterial and archaeal co-occurrence that contained two clusters, one with several *Thaumarchaeota, Verrucomicrobia* and *Actinobacteria* SVs, the other mainly with *Acidobacteria* subgroup 6 SVs. Three SVs, which could represent keystone taxa, were found to connect these clusters. If these putative keystone SVs are abundant in an aggregate, we can expect that members of both clusters are present there. The two major clusters present in the network may provide complementary functions in the soil ecosystem. There are indications that *Acidobacteria* subgroup 6, dominating one of the clusters, prefer agricultural soils with low nitrogen input where it could be involved in the slower turnover of soil organic carbon (SOC) originating from microbial necromass or plant material (Hester et al., 2018; D. D. Li et al., 2018; Navarrete et al., 2013). The soil of this study originated from a long-term nitrogen-depleted agricultural soil, thus supporting the preference to low nitrogen concentrations and SOC turnover. The other cluster included several abundant SVs from phylum *Thaumarchaeota*, which is known to be a strong contributor to ammonium oxidation in agricultural soils (Leininger et al., 2006). Compared to ammonium-oxidizing bacteria, *Thaumarcheaota* are thought to be adapted to lower nitrogen concentrations (Pester, Schleper, & Wagner, 2011), thus the nitrogen-depleted soil of this study is likely a favorable environment for them. The two clusters in our aggregate co-occurrence network could represent two distinct types of metabolism adapted to a nitrogen depleted soil: a chemoorganotroph that oxidizes SOC, and a chemolithotroph that oxidizes ammonia produced for example by ammonification from crop residues. The presence of the less abundant *Verrucomicrobia* and *Actinobacteria* SVs within the *Thaumarchaota* dominated cluster is possibly linked to an oligotrophic lifestyle (Bergmann et al., 2011; Fierer, Bradford, & Jackson, 2007), but considering the limited information that 16S rRNA gene analyses can provide for these phyla, this remains yet only a hypothesis. Shotgun sequencing and metagenomic analysis of DNA extracted from individual soil aggregates could shed more light on the nature of the associations we detected in the co-occurrence network. In general, such aggregate-level analyses of the soil microbiota, which we call “aggregatomics”, could inspire new ways of linking structure to function in soil microbial communities.

While the spatial scale that we reached in this study is not yet fine enough to reveal most microbial interactions as they may occur in microaggregates (Raynaud & Nunan, 2014), it should be able to support the development of hypotheses and experiments to understand the patterns and processes shaping the assembly of soil microbial communities and modelling their behavior (Faust & Raes, 2012; Tecon & Or, 2017). Developing DNA extraction protocols from even smaller soils samples, approaching the microaggregate-level, should be a way forward to fuel soil aggregate-oriented research (“Aggregatomics”; https://www.thuenen.de/en/bd/fields-of-activity/feld-und-laborstudien/microbiology-and-molecular-ecology/soil-aggregatomics/) for unveiling hidden patterns of functions and ecological interactions.

## Acknowledgements

We thank Ines Merbach, Department Biozönoseforschung, Helmholtz-Zentrum für Umweltforschung – UFZ, Bad Lauchstädt, Germany, for providing the soil of this study. We thank Naomie Oßwald and Jerome Rischke who supported us at the early stages of this work as bachelor students, and Karin Trescher for excellent technical assistance. The study was financially supported by the German Research Foundation (DFG; Grant number TE 383/10-1) in context of the Priority Programme “Rhizosphere spatiotemporal organization (SPP2089).

## Data accessibility statement

Sequencing data is available at the European Nucleotide Archive under the accession numbers PRJEB36881 (1^st^ experiment), PRJEB36883 (2^nd^ experiment), and PRJEB36887 (3^rd^ experiment). The qPCR data is in Supplementary file 1.

## Author contributions

Both authors contributed to the design of the experiments, analysis of the results, and writing the manuscript.

## Declaration by the authors

There are no competing financial interests in relation to the work described.

