## Supplementary information for "Hidden heterogeneity and co-occurrence networks of soil prokaryotic communities revealed at the scale of individual soil aggregates"

**Table S1** Good’s coverage index of the 16S rRNA gene sequence variants

| Soil weight class or sample type | Good’s coverage (average ± SD) |
| --- | --- |
| *1^st^ Experiment* | |
| 250 mg | 0.962 ± 0.014 |
| 125 mg | 0.965 ± 0.008 |
| 25 mg | 0.970 ± 0.009 |
| 5 mg | 0.980 ± 0.010 |
| 1 mg | 0.992 ± 0.010 |
| Control, no soil | 0.997 ± 0.001 |
| *2^nd^ Experiment* | |
| 250 mg | 0.939 ± 0.033 |
| 25 mg soil homogenate | 0.939 ± 0.031 |
| 5 mg soil homogenate | 0.945 ± 0.029 |
| 1 mg soil homogenate | 0.923 ± 0.028 |
| Control, no soil | 0.994 ± 0.005 |
| *3^rd^ Experiment* | |
| 250 mg | 0.950 ± 0.017 |
| Soil aggregate | 0.952 ± 0.032 |
| Control | 0.998 ± 0.001 |

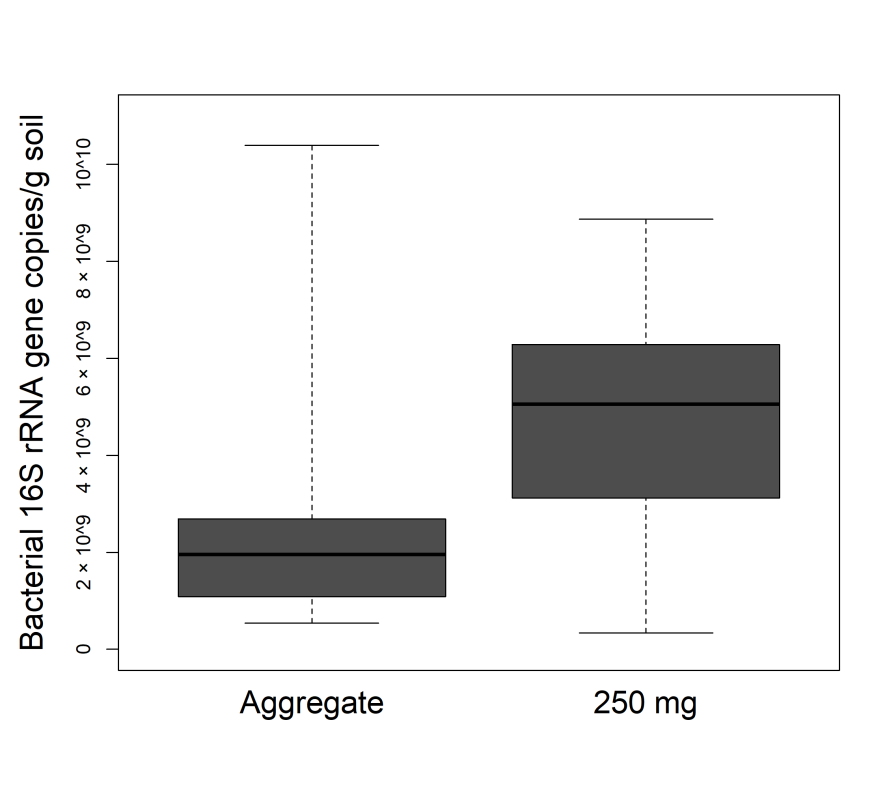

**Figure S1** Estimates of bacterial abundance in a gram of soil from the samples of the 3^rd^ experiment based on qPCR. Thick lines indicate the median values, the upper and lower hinges the 75^th^ and 25^th^ percentile, whiskers extend to the data extremes.

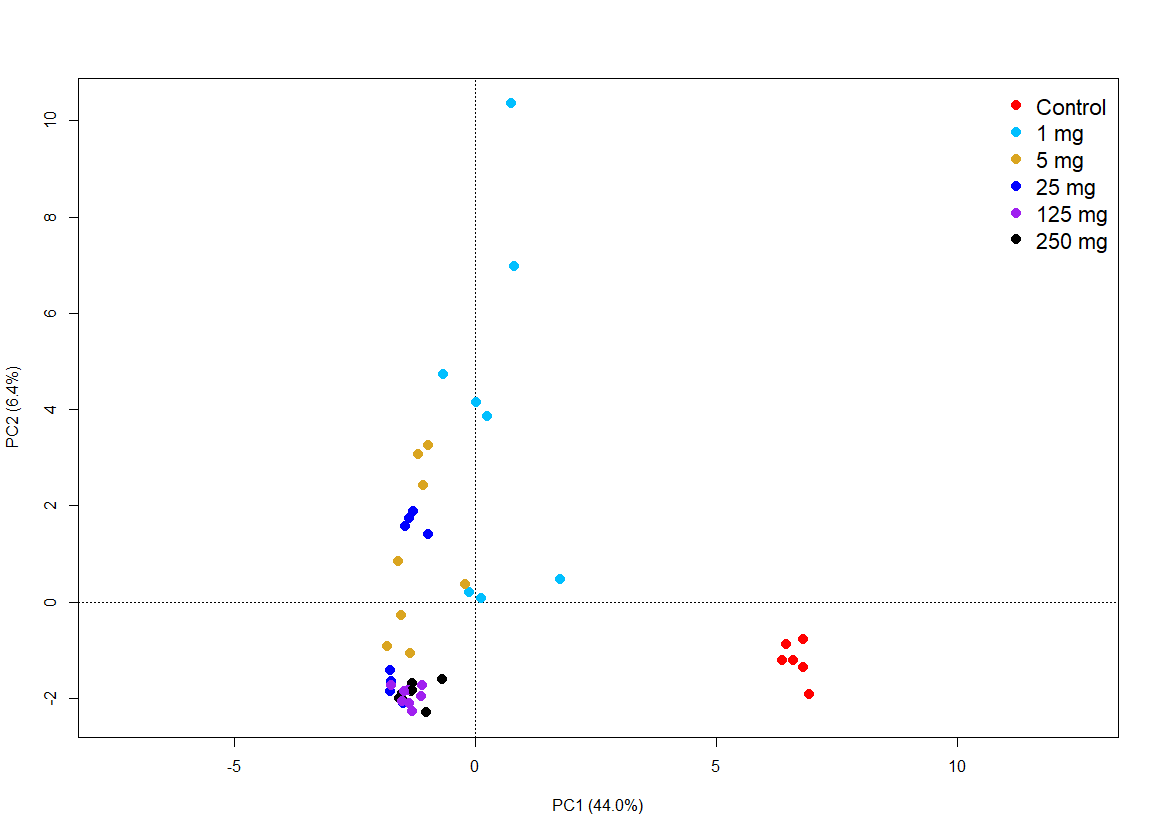

**Figure S2** Principle component analysis (PCA plot) from the 16S rRNA gene sequencing data from the 1^st^ experiment including the control samples and without the removal of potentially contaminant SVs.

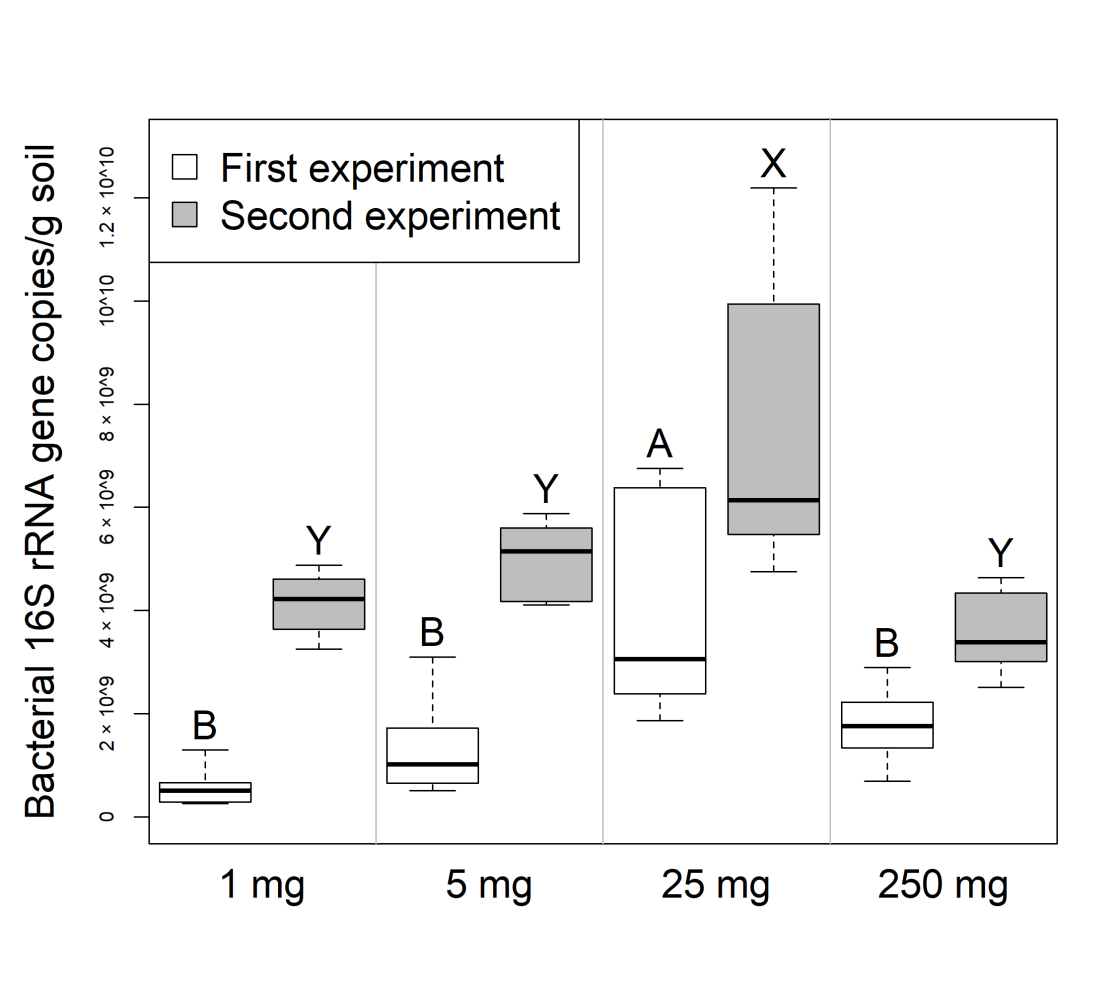

**Figure S3** Estimates of bacterial abundance in a gram of soil based on qPCR from the soil and soil homogenate samples from the 1^st^ and 2^nd^ experiments. Thick lines indicate the median values, the upper and lower hinges the 75^th^ and 25^th^ percentile, whiskers extend to the data extremes. Sample groups from the same experiment not labelled with the same letter were significantly different in Tukey’s HSD tests.
